# ChironRNA: Steric Clashes Resolution in RNA Structures via E(3)-Equivariant Diffusion

**DOI:** 10.64898/2026.03.18.712772

**Authors:** Jingyi Li, Jian Wang, Nikolay V. Dokholyan

## Abstract

Due to the limited resolution of experimental data, many determined RNA structures contain physically implausible geometries, such as severe steric clashes and missing atoms. Resolving these defects during RNA structure refinement remains a fundamental challenge. Structure dictates the function, so the geometric accuracy of RNA structure is critical for understanding biological mechanisms. However, traditional algorithms for correction have limitations because of the complexity of RNA structures. We propose ChironRNA, an all-atom diffusion model with E(3)-equivariant graph neural networks to perform RNA refinement by resolving steric clashes and completing missing atoms. In ChironRNA, we adopt a hierarchical approach, including both an all-atom diffusion model and a coarse-grained diffusion model where each nucleotide is represented by a five-point representation. Our pipeline consists of two stages: a training stage and a generation stage. The diffusion model regenerates clashing nucleotide atoms step by step by removing the noise predicted by EGNN. ChironRNA achieves an 80% clash reduction on more than 80% of the test set. It performs better on structures of less than 200 nucleotides, resulting in a high percentage of cases having over 80% clash reduction rate and 100% atom reconstruction rate. Our results demonstrate that ChironRNA successfully resolves steric clashes and rebuilds missing atoms with high precision, offering a robust solution where traditional fine-tuning or enumerative approaches fail.

## Introduction

Due to the limited resolution of experimental data, many determined RNA structures contain physically implausible geometries, such as severe steric clashes and missing atoms. Resolving severe steric clashes during RNA structure refinement remains a fundamental challenge. These geometric violations represent a high-energy state, which is thermodynamically unstable and not physically plausible^1^. RNA is involved in interactions with ligands, proteins, and other nucleic acids, thereby driving diverse biological processes such as gene expression, splicing, and translation, drug binding, ribozyme catalysis, and aptamer recognition^2,3^. Structure dictates the function, so the geometric accuracy of RNA structure is critical for understanding biological mechanisms. Consequently, the determination of accurate, clash-free 3D structures of RNA is of great importance.

There are three primary experimental ways to elucidate the structure of a protein or RNA molecule: cryo-electron microscopy^4^, X-ray crystallography^5^, and nuclear magnetic resonance^6^. Researchers have developed many computational pipelines for building protein or RNA models from experimental data^7–14^. Despite the availability of these pipelines, many experimentally determined structures in the Protein Data Bank^15^ still contain conformational defects, including steric clashes^1^, and missing atoms. These geometrical defects originate from the limitation of experimental resolution, especially at the low-resolution structures^16^. Given that severe steric clashes make a structure thermodynamically unstable, quantitative criteria have been developed to detect clashes. Van der Waals (VDW) repulsive energy can be calculated using the Lennard-Jones potential^17^. Clashes can be defined as any pairwise VDW repulsive energy exceeding an energy-based cutoff. Steric clashes are also defined by energy. Chiron^1^ defines steric clashes in protein structures as atomic overlaps that result in VDW repulsion energy greater than 0.5*K*_*b*_*T* without any covalent interactions, a disulfide bond or a hydrogen bond formed. Computing pairwise energy, especially for large molecules or molecular systems, is computationally intensive. Distance-based approaches, which define clashes based on geometric criteria, are more feasible. A clash occurs when the distance between two atoms is less than the sum of two VDW distances minus the defined cutoff. The setting of the cutoff is empirical, which could be in the range of 0.4-0.6 Å^18^. MolProbity^18^ and PROCHECK^19^ use geometrical criteria to define clashes. MolProbity provides the tool REDUCE to add hydrogen and perform optimization that consider chemical environment. The PROBE algorithm from MolProbity can perform all-atom contact analysis, calculate the clash region, and compute the clash score, which is defined as the number of serious steric overlaps per 1000 atoms. Protein Data Bank incorporates the clash score as an indicator of regions within the structures that possess unfavorable potential energy.

Despite significant progress in computational RNA or protein refinement^1,7,8,16,20–25^, current methods face limitations on structures with severe steric clashes or missing atoms. Existing approaches generally can be classified into three categories, each with distinct constraints. The energy minimization and MD-based methods, including PHENIX^8^, MDFF^20^, CNS^7^, and QRNAS^21^, primarily utilize gradient descent, simulated annealing, or molecular dynamics simulation to fine-tune the initial structure toward a conformation with improved geometry. However, when the structures contain severe steric clashes, the structural modification becomes trapped in a local minimum because the high-energy barriers prevent the conformation from fine-tuning, which makes these methods highly dependent on the initial RNA conformations. Enumerative and sampling-based methods include ERRASER^16^, RCrane^22^, and RNABC^23^. ERRASER^16^ uses an exhaustive search of nucleotide or fragment conformational space, leveraging a scoring criterion, which is the Rosetta energy score, to determine the optimal structure. RNABC utilizes forward kinematics to explore and rebuild the RNA backbone conformation. RNABC reduces the search space by anchoring the coordinates of phosphate groups and nucleobases, thereby reducing the degrees of freedom for atoms. This approach can introduce errors if the anchored atoms are misplaced^23^. The exhaustive search methods used in ERRASER face high computation costs because of the high number of possible conformations for every nucleotide. If the interaction s between nucleotides are taken into account, the computational cost increases exponentially for RNA molecules. Diffusion-based methods, including STRAND^24^ and DiffDock-P^25^, leverage a diffusion model to generate structures. STRAND is primarily designed for RNA-protein complex docking refinement. It operates on coarse-grained models on the nucleotide level, lacking the capability for all-atom refinement. Chiron^1^ utilizes discrete molecular dynamics (DMD)^26^ to perform all-atom refinement with lower computational cost compared to traditional molecular dynamics simulations. However, it is designed for proteins and cannot be directly applied to RNA. In summary, fine-tuning methods are prone to getting trapped in local minima, enumerative methods are computationally prohibitive for large-scale corrections, and existing generative models lack all-atom precision for RNA refinement. Therefore, there is a need for an all-atom level approach capable of regenerating substructures of RNA to perform refinement.

To address these issues, we developed ChironRNA, an all-atom diffusion model capable of regenerating RNA substructures to resolve severe steric clashes and complete missing atoms. Instead of relying on the initial conformation and fine-tuning, ChironRNA treats refinement as a conditional generation process. Leveraging high-quality structural regions as constraints, ChironRNA regenerates the distorted areas while maintaining the overall structure. We employ an E(3)-equivariant graph neural network (EGNN). This approach ensures rotational and translational invariance while offering a robust solution to secure physically plausible RNA conformations.

## Results

### Framework Overview

We present an equivariant diffusion-based framework for RNA structure refinement, designed to resolve steric clashes in three-dimensional RNA conformations and to reconstruct missing atoms (Figure 1). The framework utilizes conditional generation via a diffusion model to regenerate molecular substructures, thereby resolving steric clashes and reconstructing missing atoms in RNA structures. The core algorithm for generation is the Denoising Diffusion Probabilistic Model (DDPM)^27^. We use the E(3)-equivariant graph neural network (EGNN)^28^ to perform noise predictions. The EGNN respects the rotational and translational symmetries inherent to molecular structures.

**Figure 1.**
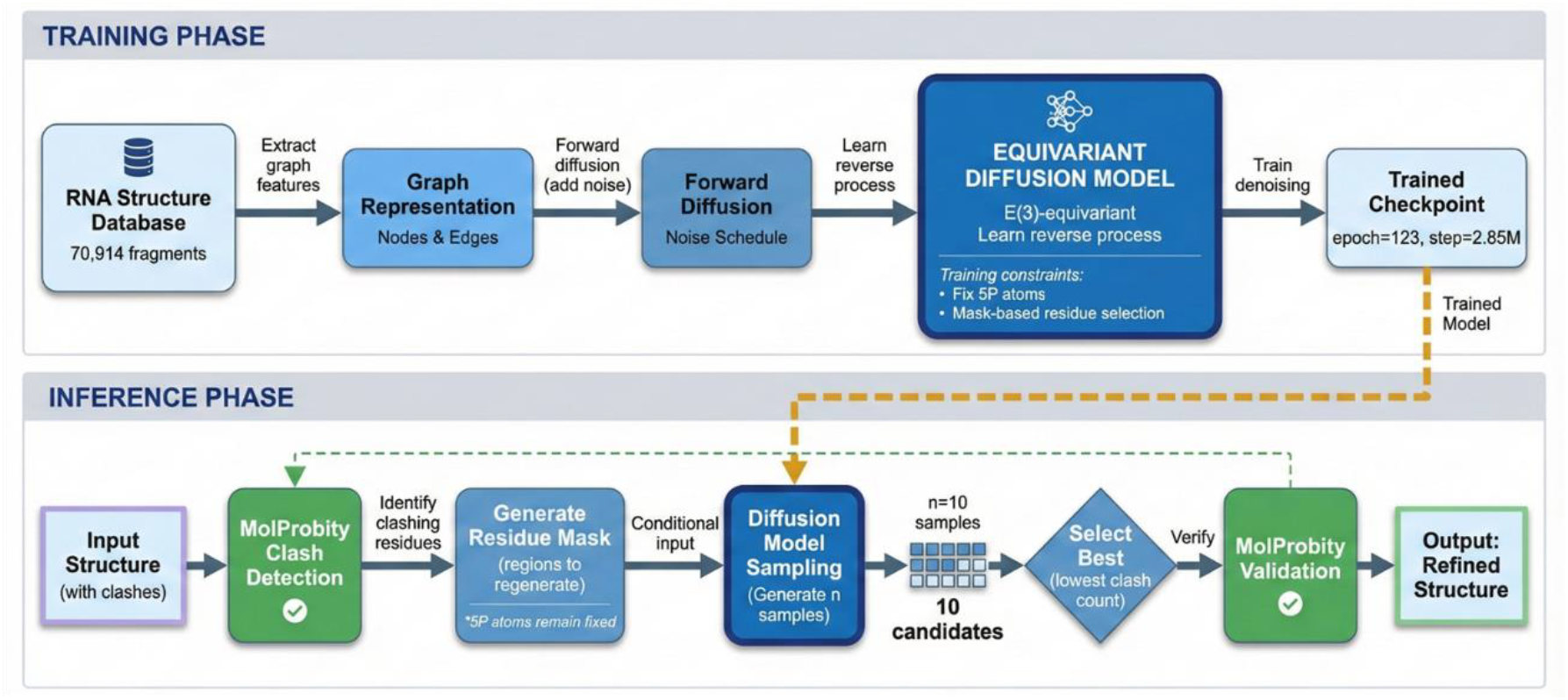
Architecture of the conditional diffusion model for RNA refinement. The diffusion model consists of a forward process and a reverse process. In the forward process, Gaussian noise is gradually added to the atomic coordinates. The EGNN is trained to learn the structural patterns by predicting the noise added at each step. In the reverse process, starting from noise, the diffusion model iteratively denoises the latent structure using the noise estimated by the EGNN. Finally, the generated candidates are evaluated by MolProbity, and the structure with the lowest clash score is selected as the final output.

The overall pipeline consists of two major components: a training process and an inference process. The training process begins with data preparation of 3D RNA structures and RNA-protein complexes from the Protein Data Bank, as well as motifs extracted from these structures. Then, the motifs and RNA structures are encoded into graph representations. Finally, these graph representations are used to train the EGNN to perform noise prediction. During the inference process, we leverage MolProbity to detect nucleotides with steric clashes. The atoms in the region with clashes are regenerated using the conditional diffusion model, conditioned on the atoms in the region with no steric clashes and partial atomic information within the clashing regions. Numerous candidates are generated, and the sample with the fewest steric clashes is selected as the output.

### Quantitative Performance Analysis

We developed a pipeline to automatically resolve clashes in RNA or its structural motifs. We built a list of RNA and motifs with steric clashes from the test dataset, consisting of 100 structures with steric clashes. We define the reduction rate as

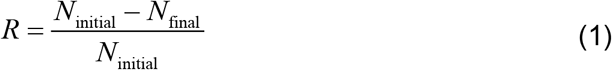

The universal high performance across clash count ranges indicates robust generalization of the model. As shown in Figure 2a, the majority of data points significantly deviate from the identity line, showing the systematic reduction with the clash resolution pipeline. Quantitatively, 60% of the tested structures have an improvement rate of more than 80%. The pipeline outperforms the structure of less than 200 nucleotides, resulting in a great percentage of cases having 100% improvement rate.

**Figure 2.**
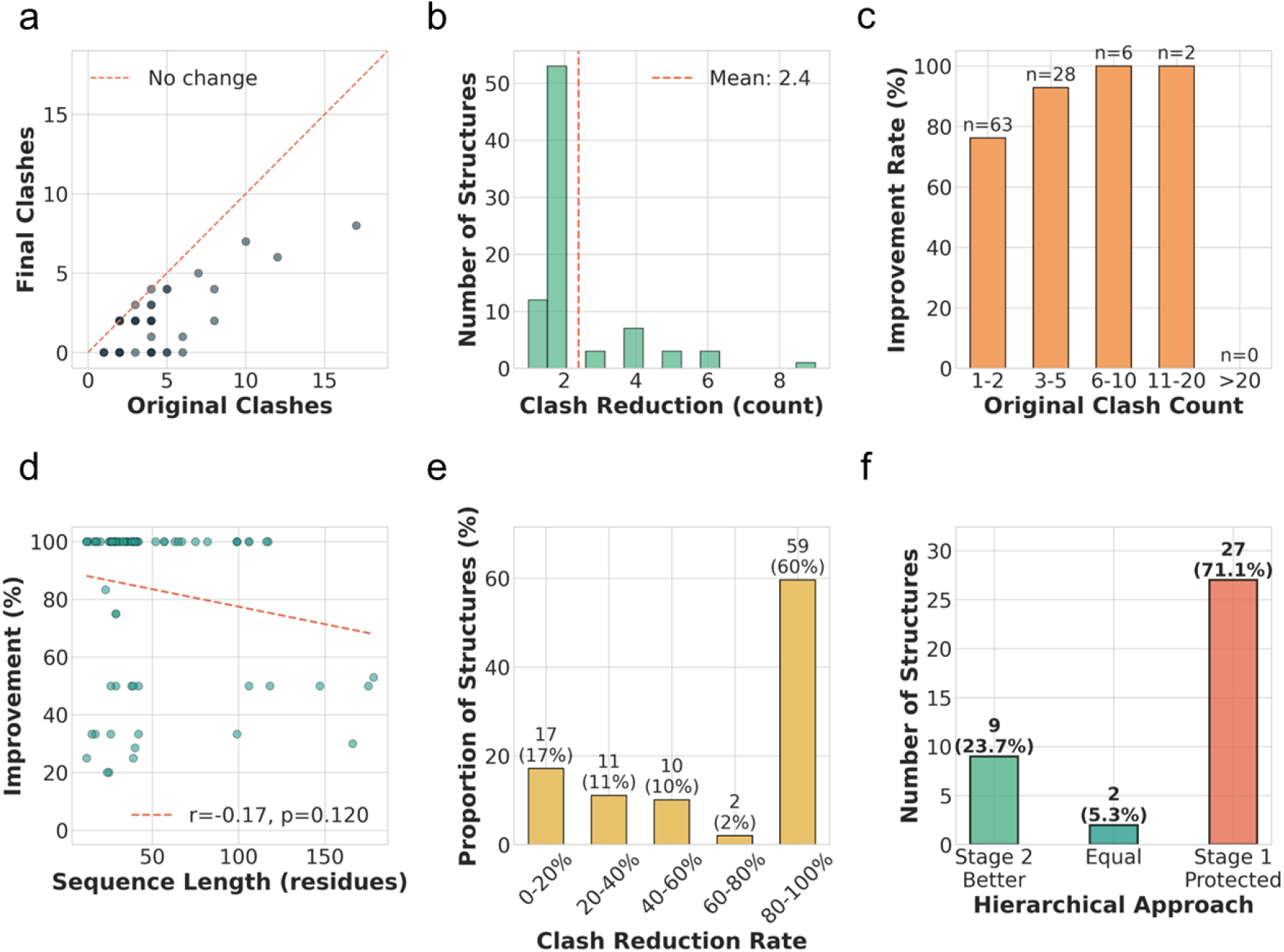
Comprehensive performance analysis of clash resolution using all-atom diffusion models. (a) Comparison of the original number of nucleotides with clashes versus the final nucleotides with clash counts after clash resolution pipeline processing across structures. Each point represents one or more structures. The darker the points, the more data points they include. Points below the red dashed line indicate successful resolution. (b) Distribution of absolute clash reduction counts across the successfully improved structures (n=82). The histogram reveals a distribution with a peak at 2, representing the most common improvement magnitude. The distribution is heavily concentrated in the 1-3 clash reduction range. The red dashed line marks the mean reduction value. (c) Improvement rate stratified by initial clash count categories, ranging from 1 to 20. The analysis demonstrates consistently high improvement rates across all ranges. 62 structures with 1-2 initial clashes have an 80% overall improvement rate**, while structures with 3 or more clashes achieved a 100% improvement rate.** (d) Correlation analysis between improvement percentage and RNA sequence length. The improvement rate is negatively correlated with the sequence length, as shown with r=-0.17 and p=0.12. (e) The distribution of clash reduction rates for all structures in the selected test dataset. 60% have an over 80% reduction rate, 17% have no more than 17% reduction rate, and 23% have a moderate success rate ranging from 20% to 80%. (f) The improvement rate for hierarchical approaches on hard cases for the all-atom diffusion pipeline. 23.7% have further improvement on the hard cases, while 74.3% have no improvement.

The model’s performance exhibits remarkable robustness across varying degrees of initial structural distortion. While the primary mode of improvement involved more than 10 severe localized steric conflicts, the pipeline maintained exceptional efficacy. This is evidenced by the 100% improvement rate observed for structures containing between 3 and 20 initial clashes (Figure 2c), suggesting that the diffusion process effectively navigates complex energy landscapes to find a clash-free local minimum. Furthermore, we observe that sequence length and improvement rate have a non-significant negative correlation (Figure 2d), indicating that the model maintains robust performance even as RNA structural complexity increases. In conclusion, these results demonstrate that the all-atom diffusion pipeline is capable of resolving steric clashes and ensuring the physically plausible generated RNA structures.

The distribution of the clash reduction rate shows the achievement of the all-atom diffusion model. However, we found that 17% of the cases were not successful, yielding a success rate of no more than 20% within this subset. This bottleneck inspired us to rethink the limitations of our constraint strategy. We hypothesized that the strict geometric constraints might be the reason for the limitations in local refinement. Specifically, having constraints on the 5-point atoms imposes limitations on the degrees of freedom for local refinement. This lack of degrees of freedom might be the origin of steric clashes. This lack leads to constraints on the possible generated structure space. We introduced a hierarchical approach to break this limitation. Relaxing the constraints on the non-C4’ five-point atoms might yield better performance for RNA clash resolution because the anchored atoms might be the source of these steric clashes. The hierarchical approach successfully yielded additional improvements in 23.7% of the “hard cases” where standard processing reached a plateau (Figure 2f). By decoupling these constraints, the hierarchical approach successfully resolves the steric traps induced by rigid anchoring.

### Case Study of Atomic-level Clash Resolution in RNA Structures Using Equivariant Diffusion Models

To evaluate the atomic-level performance of our diffusion pipeline, we analyzed representative RNA structures and visualized them with red dots representing steric clashes. Figure 3a shows an Internal loop from PDB 1I95 defect with a high energy state, including 3 pairs of severe steric clashes, with maximum overlap up to 0.892 Å. Atoms in different nucleotides with no Watson-Crick interaction form pairs of steric clashes. As shown in Figure 3b, the all-atom diffusion model successfully resolved the steric clashes by regenerating the atoms in the nucleotides that were involved in the steric clashes. Diffusion process guided by EGNN did not fine-tune the structure, but reconstructed the local geometry, leading the system to a state with no steric clash or missing atoms. The second case is an RNA PDB 4D8Z with 188 nucleotides, which is more complex in RNA topology. The challenge for local optimization is the interactions between nucleotides. The local conformation interacts with the whole structure. Resolving the local clashes needs structural features for the whole structure. The initial conformation contains 4 clashes, with a maximum overlap 0.677Å. Figure 3d shows that the all-atom pipeline achieves 75% success rate. The success of clash resolution shows that the diffusion model utilizes the long-range interactions captured by EGNN to refine local structures without forming new steric clashes with constraint regions. ChironRNA successfully has a balance on local atomic rearrangement and global stability constraints for the structure with 188 nucleotides. Finally, we show a highly constrained Hairpin loop from PDB 1QA6. This case shows the importance of a hierarchical strategy. Initial conformation contains clash pairs with 0.804Å overlap. With five-point atom rigid anchoring, the all-atom pipeline stagnates in the structural refinement (Figure 3f). The strict anchoring of the atomic degree of freedom makes the system trapped in a part of conformation space. However, the hierarchical approach successfully breaks the rigid coordinate constraints. By releasing the degree of freedom for non-C4 five-point atoms, the hierarchical strategy makes the system have more degrees of freedom. The expansion of degrees of freedom provides more possible conformations for ChironRNA, enabling the success of the hierarchical approach (Figure 3g). This strongly proves that hierarchical special sampling can overcome the local minimum in the single-stage optimization methods.

**Figure 3.**
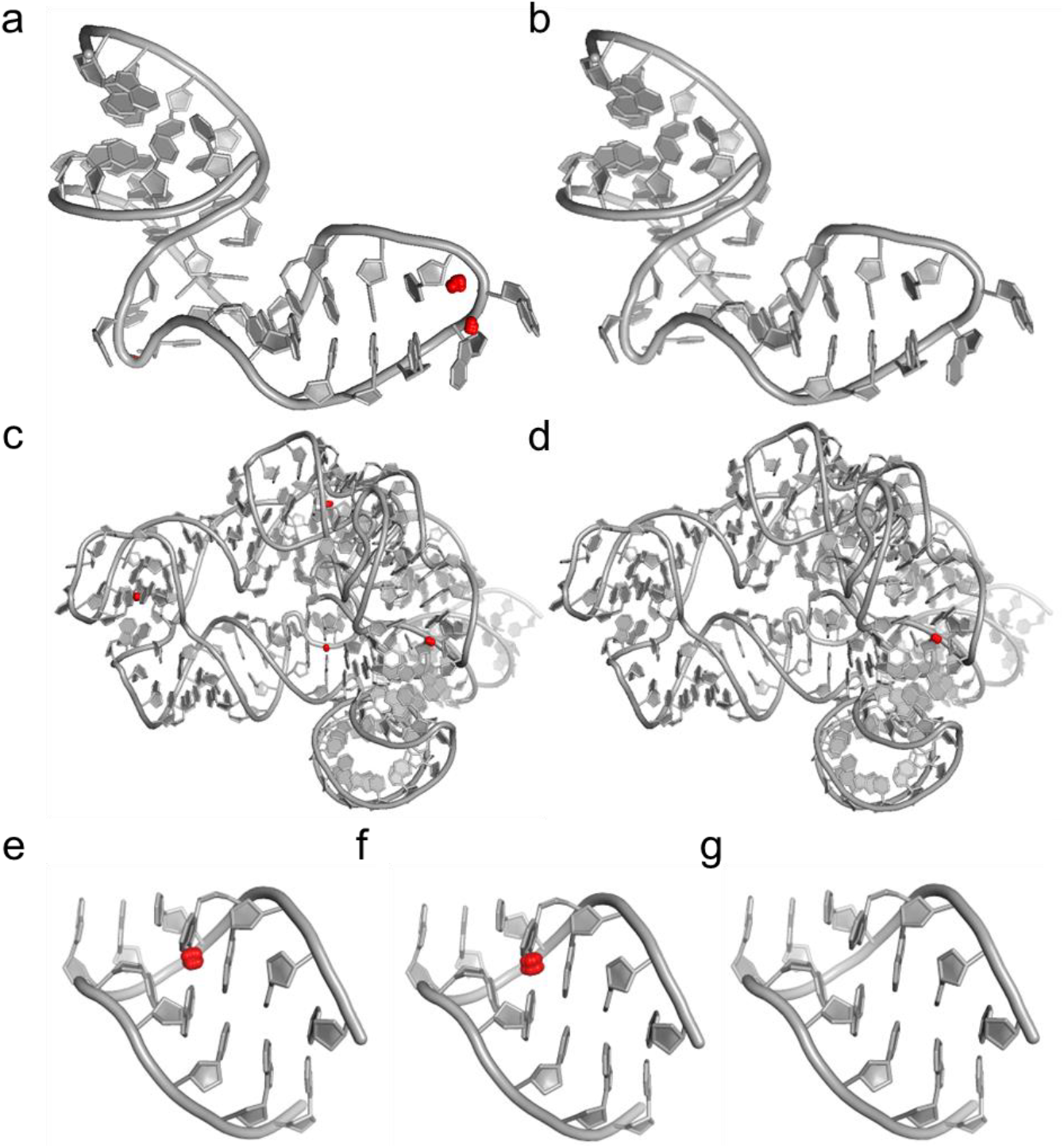
Case study of Atomic-level clash resolution in RNA structures using equivariant diffusion models. (a-b) An example of complete clash elimination in a 35-residue structure. (a) is the original structure with 3 clash pairs with the worst overlap of 0.892 Å, while (b) shows the output without clashes (0 clashes).(c-d) An example of a large RNA motif structure with 188 nucleotides. (c) The original structure contains 4 clash pairs labeled by the red dots, with the worst overlap of 0.677 Å. (d) shows the output of the all-atom refinement pipeline, which reduces the clash to 1 pair with a 75% improvement. (e-g) It is an example of the capability for handling the challenging case (12 residues) that the all-atom pipeline cannot handle. (e) shows the original structure with 1 clash pair, with the worst overlap of 0.804 Å. (f) demonstrates that the improvement of the all-atom pipeline is unsuccessful for this case, failing to resolve the overlap, while the hierarchical approach (g) achieves complete success on the challenging case, outputting the structure with 0 clashes.

### Reconstruction of Missing Atoms

We developed a pipeline for fixing missing atoms for RNA or motif structures. The pipeline detected nucleotides with missing atoms and reconstructed nucleotides with other regions of structures as constraints. To evaluate the performance of the pipeline, we randomly deleted the non-five-point atoms, reconstructed the structures, and then calculated the RMSD between the initial structure and the reconstructed structures. We chose an example for visualization, as shown in Figure 4 a and b. In order to quantify the overall performance, we had the reconstruction for 90 motifs and then plotted the RMSD between the output and the initial structures. Figure 4 c shows the high accuracy of the reconstruction pipeline, demonstrating the capability to restore chemically accurate atomic coordinates for structures with missing atoms while maintaining structural integrity.

**Figure 4.**
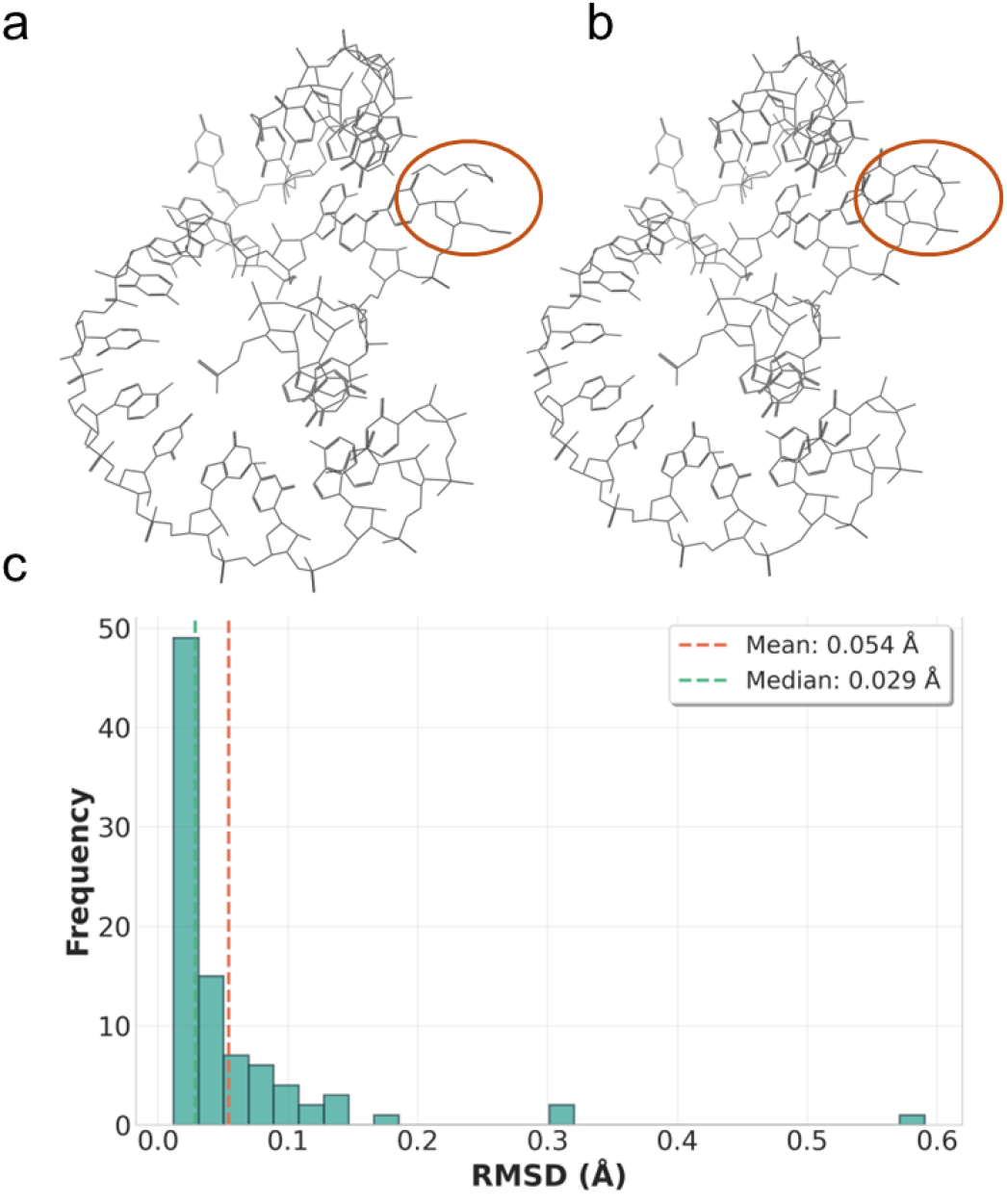
Atom missing recovery performance using the all-atom diffusion model. (a) Representative example is an internal loop from the PDB database 6WNT with deleted non-five-point atoms. The motif has 8 atoms missing from one residue, as shown in the red circle, including backbone atoms (OP2, O5′, C5′, O2′, O3′) and base atoms (O2, C4, C6), representing typical incomplete structural data scenarios. (b) Recovered structure with all-atom diffusion mode. The model successfully reconstructed all missing atoms with high accuracy, and the RMSD between the reconstructed structure and the initial structure from PDB is 0.018 Å. (c) Distribution of all-atom RMSD values across the validation dataset (n=90 successfully recovered from 100 test cases). The histogram shows highly accurate recovery performance with a median RMSD of 0.029 Å, labeled by the green line, and a mean RMSD of 0.054 Å, labeled by the orange line, indicating sub-angstrom precision across diverse RNA structural motifs for atomic reconstruction for the majority of structures.

Finally, we performed a visualization of the all-atom diffusion process of a stem consisting of 6 nucleotides. The initial conformation is Gaussian noise with all the atoms crowded in a close region. During the inverse diffusion process, the atomic RNA structure is generated by removing the clashes. Figure 5 provides a visualization of clustering the atoms from different nucleotides, forming the backbone shape, and adjusting the atomic coordinates.

**Figure 5.**
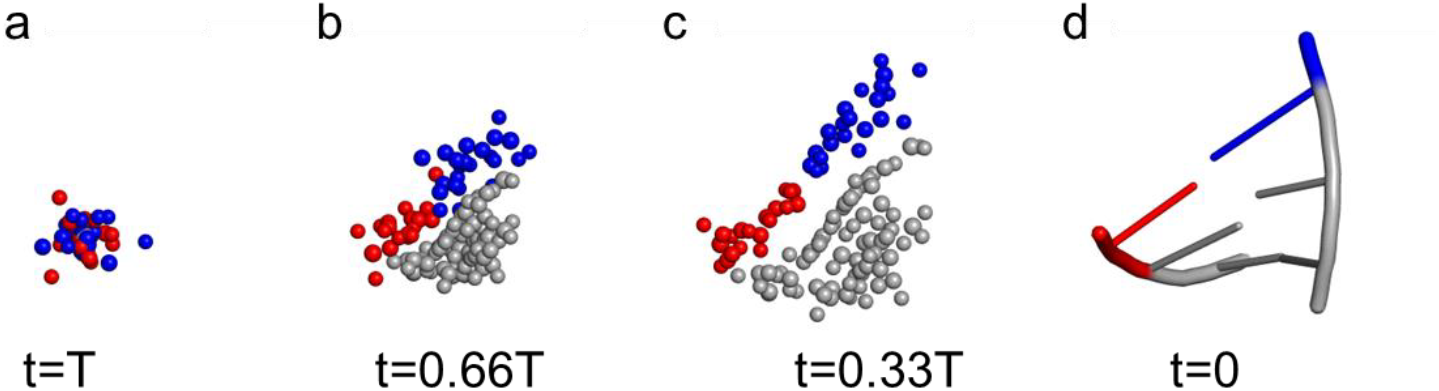
Reverse diffusion process in all-atom RNA clash resolution. (a) The initial conformation of a stem consisting of 6 nucleotides. Different colors represent different nucleotides. The atoms with gray represent the constraint atoms, which recover the input conformation. The gray and red represent the atoms in different nucleotides to be generated. (b) After the diffusion process, atoms from different nucleotides disperse, clustering into different groups. (c) Atoms move and form a shape like the backbone conformation. (d) After adjusting the atomic coordinate, the final conformation forms.

### Diffusion Process Visualization

To elucidate the underlying mechanism of the conditional generation in the RNA refinement process, we show the reverse diffusion process with a representative RNA motif structure. This process reveals the generative process of an RNA motif using a diffusion model with an EGNN from pure noise. In this demo, we track the 3D coordinates of a stem with 6 nucleotides during the RNA refinement process. As shown in Figure 5, the gray spheres represent constraint atoms, representing the region with high-fidelity substructures. Conversely, the red and blue spheres represent the region with structure distortion, which needs to be regenerated. In this region, adding Gaussian noise destroys the substructure, leaving the atoms crowded in a narrow space in highly disorder state The forward diffusion process destroys the structures, removing all the initial poor geometry. During the reverse diffusion process, the EGNN guides the model to separate the overlapping atoms. The diffusion model gradually clusters the atoms belonging to the same nucleotide, forming different groups, as shown in Figure 5b. Based on the atomic features, the diffusion model efficiently clusters the atoms. Then, the clustered atoms form physically plausible structures. In this stage, the initial backbone of the RNA structure is formed, as shown in Figure 5c. At the final stage of the generation process, the diffusion model fine-tunes the atomic coordinates, forming a physically plausible structure by incorporating the interactions between atoms within and across the nucleotides. With the geometry features captured by the EGNN, the diffusion model regenerates the substructures, fixing all missing atoms and reducing the steric clashes. The final structure is shown in Figure 5d. This process reveals the capability of atomic regeneration with partial constraints.

## Discussion

We have proposed ChironRNA, an EGNN-based all-atom diffusion model for RNA steric clash resolution, which is a challenging problem for RNA 3D structures. The results show that the diffusion pipeline is capable of solving severe steric clashes and the missing atom problem. Different from the traditional gradient descent, MD simulation, and exhaustive sampling, diffusion models regenerate RNA substructures to perform RNA refinement, avoiding the dependency on the initial conformation. Our approach achieves atom-level refinement, maintaining the consistency of the global structure.

We found that corrupting the local structure and then regenerating the local structure is an efficient strategy to perform RNA refinement. Some of the traditional approaches, including the MD and gradient descent approach, are primarily fine-tuning to the initial conformation. However, when the initial structures contain a high energy barrier to overcome, the conformations tend to get trapped in the local minima, resulting in the failure of the refinement. By corrupting the local structure, we destroyed the high-energy barrier during the noising process. During the denoising process, the diffusion model utilizes the local environment as conditioning and guides the atoms to the physically plausible conformations. For the hard cases, we found that the hierarchical approach has further improvement. Anchoring the five-point atom constrains the degree of freedom; the misplacement of the five-point atoms caused the overall structure defect. We design a hybrid masking strategy, which captures not only neighbors but also long-range interactions, which is crucial for capturing the complex RNA folding patterns.

Comparison with the existing methods, traditional approaches, including PHENIX^8^ and RNABC^23^, utilize the gradient descent or forward kinematics. Despite the performance for the fine-tuning tasks, they are not capable of fixing severe distortions and atom missing, while our diffusion-based approach is not highly dependent on initial conformation. Compared to the enumerative approach, including ERRASER^16^ it has high accuracy in rebuilding single nucleotide. However, it rebuilds the structure one by one, ignoring the long-range interaction between nucleotides. Graph representation of the graph and message passing enables the interaction between nucleotides. Diffusion model-based approach^24 31^ aims for a different task, which is RNA-protein docking and molecular design in protein pockets instead of RNA refinement.

We hypothesize that the coordinate of the C4’ atom is accurate and treat the atom as the conditioning for refinement. However, if the large deviations happen, the refinement might fail. Because of the EGNN and VRAM constraints, we set the maximum length to 200, which is not capable of handling larger RNA structures. At the same time, we found that the refinement success rate was negatively corrected with sequence length. We did not use the full connection network, so we cannot capture all long-range interactions. We lack direct experimental result constraints, resulting in the output conformation may be physically plausible but not have good alignment with the experimental result.

Graph construction is facing the dilemma of precision and efficiency. A full-connection graph has high precision and efficient message passing, while fewer connections use less VRAM, and it is computationally efficient. For processing longer RNA structures, we will test the performance of sparse graph encoding, including a global virtual node, which acts as the virtual node that connects all-atoms for one nucleotide^32^, or K-nearest-neighbors^33^, finding a balance between precision and efficiency for RNA graph representation. Inspired by Ding and Dokholyan^34^, who perform RNA structure prediction with discrete dynamics simulation, incorporating hydroxyl radical probing data as physical constraints, our future work aims to develop the diffusion model with guidance, incorporating experimental data. Alternatively, future models could incorporate experimental data, such as cryo-EM density maps, directly into graph representations.

## Methods

### Data Retrieval and Curation

The training dataset is constructed from experimentally determined RNA structures in the RCSB Protein Data Bank^15^. RNA, RNA-Protein, and RNA-DNA complexes are retrieved from the PDB database via Application Programming Interface (API), with each successful retrieval stored in a PostgreSQL database table that saves complete RNA structures.

Quality control filters were developed to ensure structural integrity. Structures are retained only if they contain standard RNA nucleotides (Adenine, Uracil, Guanine, and Cytosine), while entries with non-standard modifications or incomplete coordinate information are excluded. The cleaning process removes all non-RNA components, including associated proteins, water molecules, ions, and ligands, yielding RNA structures suitable for training.

### Secondary Structure Annotation

Secondary structures of the RNA structures are computed using MC-Annotate^29^, which analyzes the 3D coordinates to identify inter-nucleotide interactions, outputting the Watson-Crick (WC) interactions with dot bracket notation. Secondary structures are used to derive motifs and graph representations for RNA structures.

### Structural Fragment Extraction

RNA motifs are recurrent substructures within RNA structures. To increase training data diversity and learn the features of RNA motifs, we extracted and stored RNA motifs, including stems, hairpin loops, internal loops, and multi-way junctions, by decomposing RNA structures. Since we strictly require WC interactions for these motifs, we use secondary structures for extraction, which is computationally more efficient compared to the graph isomorphism algorithm used for non-WC motif extraction.

By identifying successive structural units through topological information, the fragment extraction algorithm processes the secondary structure tree to identify motifs. Hairpin loops are extracted as terminal elements containing a stem capped by an unpaired loop region. Internal loops and bulges are stems with an unpaired region inserted in the middle. Junction elements are extracted at branch points where three or more stems extend from the convergence point. The extraction pipeline populates a motif table containing over 70,000 entities in the database. Each record stores the fragment type, sequence, secondary structure annotation, and 3D atomic coordinate representations. The motif table provides a more diverse and abundant dataset compared to full-length RNA structures. The dataset is then partitioned into training, validation, and test sets with a ratio of 80%, 15%, and 5%, respectively. We used clash-free structures for the training and validation datasets.

### Graph Representation of RNA Structures

Encoding the RNA and motif structures is the process of encoding the PDB data into a different representation, which facilitates the incorporation of RNA structural features. Graph representations are naturally suitable for molecular representation s^30^ because we can treat the nucleotides or atoms as discrete nodes connected by edges that represent interactions.

Graph representation of RNA structures can be hierarchical: residue-based, five-point coarse-grained, and all-atom based, which include all heavy atoms. In a residue-based graph, C4’ atom represents a nucleotide. In a five-point atom-based graph, a five-point atom represents a nucleotide. We represent the RNA structure as a geometric graph *G* = (*V, E*), where nodes *v*_*i*_ ∈*V* correspond to individual heavy atoms, and edges *e*_*ij*_ ∈ *E* encode structural and spatial interactions. The encoding process constructs graphs with PDB files, including atom index, atom type, nucleotide type, and atomic coordinates.

RNA structures are encoded as attributed graphs where nodes represent heavy atoms and edges encode structural interactions. The graph construction algorithm parses the PDB files to extract the atomic coordinates and chemical characteristics, and encodes them into a graph format, keeping the structural information.

### Node Feature Construction

Node features are constructed as concatenated vectors encoding both chemical identity and sequential position, including (1) residue type one-hot encoding, (2) atom type one-hot encoding distinguishing the 85 unique heavy atoms defined in the PDB standard, and (3) sinusoidal positional encoding to capture the nucleotide sequence information. In all-atom representations, the residue one-hot is based on the residue type: A, U, G, and C. We use a unique index for each atom based not only on its atom type but also on its residue position, capturing position information on an atomic level compared to encoding solely on atom type. The positional encoding, illustrated in Equations (2) and (3), encodes the position information of each residue, providing the graph with information about the sequential location of each residue within the RNA chain. We use a sinusoidal scheme where each position is represented by sine and cosine functions at different frequencies.

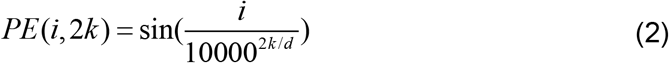

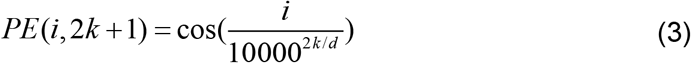

The final node feature vector is obtained by concatenating the feature vectors. This design allows the network to distinguish between chemically identical atoms that occur in different sequential contexts.

### Edge Construction and Features

For all-atom representation, we use one-hot encoding for edge features, containing four types. Intra-residue edges build the connection atoms within one residue, capturing the local geometry constraints and covalent bonding. Sequential edges connect all-atoms between adjacent nucleotides, representing the backbone connectivity and the geometric interactions between atoms in adjacent nucleotides. Base-pair edges connect atoms in the nucleobases of residues engaged in Watson-Crick interactions, as identified from secondary structure annotation. Spatial edges connect the atoms in the residues that are geometrically close to each other. We use the threshold of C4’ atom distance less than 8Å, capturing the nucleotides which are geometrically close but without WC interactions or covalent bonds.

We have five different types of edge encoding, which are C4’-C4’ interactions of adjacent residues, C4’-C4’ interactions of non-adjacent residues, non-C4’ atom with C4’ atom within the same residue, non-C4’ atom pairs within the same residue, and non-C4’ atoms between residues whose C4’ distance is less than 14Å.

### Diffusion Model with E(3) Equivariant Graph Neural Network

Diffusion models are inspired by the diffusion process, simulating the diffusion process by adding Gaussian noise to the normalized data. Then, perform the generation process by removing the noise, which is usually learned by a neural network. We use a hierarchical architecture, consisting of base building blocks, composite blocks, complete neural networks, and diffusion wrappers. Base building blocks include a graph convolutional layer (GCL), handling node feature updates, and an equivariant layer, handling coordinate updates following E (3) equivalent, which is invariance to rotations and translations. Composite blocks combine multiple GCL layers with one equivariant layer. The EGNN network stacks multiple equivariant blocks with embedding layers. Diffusion model wraps EGNN with time conditioning, performing the denoising process for RNA structure generation.

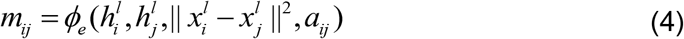

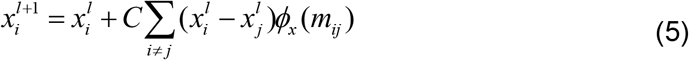

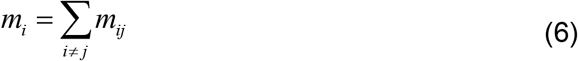

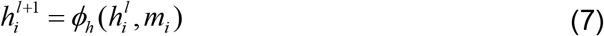

As shown in Equation (4), the network ensures translation invariance by utilizing the relative squared distance as input rather than absolute atomic positions. The node embeddings *h*_*i*_ and *h*_*j*_ represent the atom feature, while *a*_*ij*_ represent interaction features. Equation (5) details the E(3)-equivariant coordinate update: atomic positions are iteratively refined using a vector field oriented along the radial direction, thereby maintaining rotational and translational symmetry. Equations (6) and (7) show the GCL feature update. The message from the neighbor is aggregated into the node and is then used to update the edge features, as shown in Equation (7). The stack of base building blocks is EGNN, which is wrapped in the diffusion process. The diffusion process consists of the forward process and the reverse process, which is a generative process.

The forward process gradually corrupts the atomic coordinates with Gaussian noise according to a variance schedule.

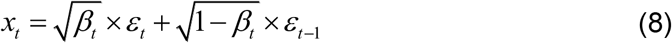

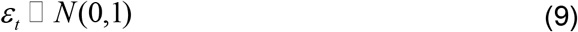

The repetitive process is simplified to a one-time sampling, where 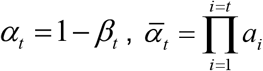. *ε* is the neural network output.

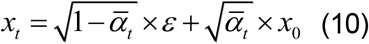

The reverse process is learned by the EGNN to iteratively denoise the structure, recovering the clean atomic coordinates from a noise distribution.

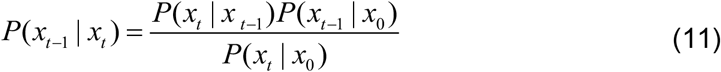

RNA structures are obtained from the denoising process step by step.

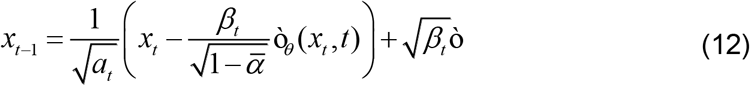

RNA structure is not dependent on the axis, which means that having translation and rotation does not change the properties of RNA molecules. The energy of RNA molecules is invariant; the relative position vector, velocity, and flux vector in the RNA are independent to the reference frame. We use the EGNN to maintain the equivalence of the RNA structures. Compared to using reshuffling on the 3D coordinate, it saves the data augmentation process by maintaining the geometric properties at the model level.

### Constrained Diffusion Training

In the inference stage, the region with clashes is regenerated, while the high-quality regions are fixed as constraints to maintain consistency with the input structure. The goal is to modify only the atoms involved in steric clashes while preserving the overall fold. The constrained generation framework employs a partial masking strategy where atoms are divided into two sets: fragment mask and linker mask.

For RNA structures, we adopt a five-point atom as a coarse-grained representation of each nucleotide. The diffusion process generates the all-atom structure with the coarse-grained representation as conditioning. The 5-point representation comprises C4’, which is the central sugar carbon, C1’, which is the glycosidic carbon, and the three base atoms that define the nucleotide base orientation: N9, C6, and C2 for A and G, and N1, N3, and C5 for U and C. The five-atom representation captures the geometric and chemical properties of each nucleotide while reducing the complexity of representations by reducing the number of graph nodes and edges.

The fragment mask defines the anchored atoms, while the linker mask defines the atoms to regenerate. The non-five-point atoms in the region with steric clashes are defined as a linker mask, while the atoms in the region with high quality and the five-point atoms in the region with steric clashes are defined as a fragment mask. Architecture enables the diffusion model to generate chemically reasonable RNA structures with constraints.

To simulate the scenarios where clashes are more prevalent, we developed a hybrid masking strategy. We employ a hybrid masking strategy to assign the atoms to nucleotides of different masking types, combining two complementary approaches for selecting nucleotides. Phase 1 implements sequence-based and secondary structure-based masking, which captures a high percentage of the training dataset. The algorithm randomly selects 1-5% of nucleotides as the initial seed for each selection. The seeds are then expanded to neighboring residues with independent 0.4 probabilities. The seeds are expanded to nucleotides, which have a base pair with them with a 0.3 probability. The expansion strategy creates sole nucleotides or contiguous regions for simulating the real steric clash region, corresponding to clashes within isolated nucleotides and clashes between nucleotides with covalent or WC interactions. Interactions occur between nucleotides that are distant in sequence but spatially proximal. Phase 2 implements 3D structure-based masking using a K-nearest pairs algorithm. The algorithm calculates the pairwise C4’ distance between all nucleotides and selects the K-nearest pairs that satisfy exclusion criteria to avoid overlapping with phase 1. The exclusion criteria exclude the third-order sequence neighbors, neighbors with direct base pairs, and the second-order neighbor of nucleotides with direct base pairs. The parameter K is computed based on a randomly sampled severity level, which is set based on the clash ratio in the RNA structure in RNA data. We set uniform distributions for different severities. We set 60% severity=0, 30% of severity is between 0 and 0.2, 9% of severity is between 0.2 and 0.5, and 1% severity is between 0.5 and 1. The number of chosen nucleotides, K, is calculated by

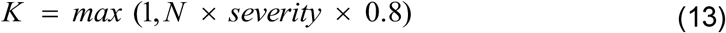

Where N is the sequence length. The nucleotide assigned with linker mask ranges from 1 pair to approximately 80% of sequence length, and 90% of the cases fall between 1 and 16% of sequence length, which is close to the statistic of the clash ratio in RNA structures.

The final mask is computed as a hybrid approach, with the union of residues selected by either phase. The hybrid approach captures the continuous regions that are likely to have clashes based on sequence, secondary, and tertiary structures. Randomness ensures the diversity of the training dataset, promoting robust learning of conditional generation, boosting the generalization of the EGNN used in the diffusion model.

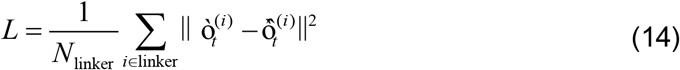

The model is optimized with a mean squared error as the loss function, aiming at the linker regions that are generated during the refinement process. The loss *L* is computed as the average square difference for atoms with a linker mask between the truth noise 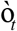 and the predicted noise 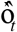 at each timestep, where *N*_linker_ denotes the total number of atoms with a linker mask. To balance computational efficiency and structural complexity, the maximum nucleotide number is set to 200. The model uses a 64-dimensional feature space as the default setting. The diffusion process has 100 timesteps, employing a polynomial noise schedule.

### Clash Detection and Resolution

We integrated the MolProbity software suite, which is the gold standard method for identifying steric clash. We utilize MolProbity to detect the region with severe steric clashes, providing information for masking definition. The detection pipeline consists of two stages: hydrogen placement and contact analysis. PDB data does not always include the hydrogen atoms, and hydrogen atoms are part of the atom-atom interaction and steric clashes, resulting in the hydrogen placement, which is an important part of the steric clash analysis^18^. Then, we integrated the probe modulus in the pipeline to analyze all pairwise atomic contacts and report the overlap values where atoms interpenetrate. The threshold we set for the distance-based overlap is 0.6Å, corresponding to severe clashes^18^.

We sample multiple candidate structures and select the best candidates based on the clash number, which is calculated by MolProbity. Given the input structure with detected clashes, the pipeline generates multiple structures (default 10) to increase the diversity of output structures, boosting the probability of obtaining an optimal candidate. For each candidate, the MolProbity analysis is repeated to quantify the clash count, and the candidate with the minimal clash number is set as the refined output.

We developed a protection mechanism to avoid degradation in difficult cases. The algorithm compares the best candidate and the input. If the best candidate has more clashes than the input, the output is returned as the input structure, preventing the model from introducing more clashes than the input structure.

We built an adaptive pipeline for solving those hard cases by regenerating the non-C4’ five-point atoms. To address the most recalcitrant cases with a success rate of no more than 60% in the all-atom refinement pipeline, we introduced a hierarchical refinement strategy. Beyond the all-atom refinement model, we built a coarse refinement strategy by generating five-point atoms with local and C4’ atom constraints, relaxing the constraints on non-C4’ five-point atoms, allowing a greater degree of freedom for atomic-level refinement. The two-stage algorithm first utilizes the five-point refinement to generate the five candidates of the five-point atoms coarse-grained model. Then, the all-atom pipeline generates 5 samples for each output of the five-point refinement algorithm. The hierarchical approach refines the structures stepwise, expanding the refinement space compared to the all-atom refinement. MolProbity is then employed to choose the optimal structure with fewer clashes.

## Data Availability

Data available on reasonable request.

## Acknowledgments

We acknowledge support from the National Institutes of Health (R35 GM134864), the National Science Foundation (2040667), and the Passan Foundation. This project was also supported by the Penn State College of Medicine’s Artificial Intelligence and Biomedical Informatics Program.

## References

1. Ramachandran, S., Kota, P., Ding, F. & Dokholyan, N. V. Automated minimization of steric clashes in protein structures. Proteins: Structure, Function and Bioinformatics 79, 261–270 (2011).

2. Mortimer, S. A., Kidwell, M. A. & Doudna, J. A. Insights into RNA structure and function from genome-wide studies. Nat. Rev. Genet. 15, 469–479 (2014).

3. Cech, T. R. & Steitz, J. A. The noncoding RNA revolution - Trashing old rules to forge new ones. Cell 157, 77–94 (2014).

4. Kühlbrandt, W. The resolution revolution. Science (1979). 343, 1443–1444 (2014).

5. Riley, R. & Chapman, V. A three-dimensional model of the myoglobin molecule obtained by X-ray analysis. Nature 181, 662–666 (1958).

6. Wüthrich, K. Protein structure determination in solution by NMR spectroscopy. Journal of Biological Chemistry 265, 22059–22062 (1990).

7. Brunger, A. T. et al. Crystallography & NMR System: A New Software Suite for Macromolecular Structure Determination. Acta Cryst 54, 905–921 (1998).

8. Adams, P. D. et al. PHENIX: A comprehensive Python-based system for macromolecular structure solution. Acta Crystallogr. D Biol. Crystallogr. 66, 213–221 (2010).

9. Glime, J. M. Auto-DRRAFTER: Automated RNA Modeling Based on Cryo-EM Density. Methods in Molecular Biology 2568, 1–18 (2022).

10. Li, T., Cao, H., He, J. & Huang, S. Y. Automated detection and de novo structure modeling of nucleic acids from cryo-EM maps. Nature Communications 15, (2024).

11. Kendrew John C. & G. Bodo. DeepTracer for fast de novo cryo-EM protein structure modeling and special studies on CoV-related complexes. Proceedings of the National Academy of Sciences 118, e2017525118.

12. Vernon, R., Shen, Y., Baker, D. & Lange, O. F. Improved chemical shift based fragment selection for CS-Rosetta using Rosetta3 fragment picker. J. Biomol. NMR 57, 117–127 (2013).

13. Wang, X., Terashi, G. & Kihara, D. CryoREAD: de novo structure modeling for nucleic acids in cryo-EM maps using deep learning. Nat. Methods 20, 1739–1747 (2023).

14. Jamali, K. et al. Automated model building and protein identification in cryo-EM maps. Nature 628, 450–457 (2024).

15. Berman, H. M. et al. The Protein Data Bank. Nucleic Acids Res. 28, 235–242 (2000).

16. Chou, F. C., Sripakdeevong, P., Dibrov, S. M., Hermann, T. & Das, R. Correcting pervasive errors in RNA crystallography through enumerative structure prediction. Nat. Methods 10, 74–76 (2013).

17. Jones, J. E. E. On the Determination o f Molecular Fields.—II. From the Equation of State of a Gas. 4, 463–477 (1851).

18. Davis, I. W. et al. MolProbity: All-atom contacts and structure validation for proteins and nucleic acids. Nucleic Acids Res. 35, (2007).

19. Laskowski Roman A. PROCHECK: a program to check the stereochemicai quality of protein structures. J. Appl. Crystallogr. 26, 283–291 (1992).

20. Trabuco, L. G., Villa, E., Mitra, K., Frank, J. & Schulten, K. Flexible Fitting of Atomic Structures into Electron Microscopy Maps Using Molecular Dynamics. Structure 16, 673–683 (2008).

21. Stasiewicz, J., Mukherjee, S., Nithin, C. & Bujnicki, J. M. QRNAS: Software tool for refinement of nucleic acid structures. BMC Struct. Biol. 19, 1–11 (2019).

22. Keating, K. S. & Pyle, A. M. RCrane: Semi-automated RNA model building. Acta Crystallogr. D Biol. Crystallogr. 68, 985–995 (2012).

23. Wang, X. et al. RNABC: Forward kinematics to reduce all-atom steric clashes in RNA backbone. J. Math. Biol. 56, 253–278 (2008).

24. Al-zeqri, M., Franke, J. K. H. & Runge, F. STRAND: Structure Refinement of RNA-Protein Complexes via Diffusion. bioRxiv 2025.06.30.662415 (2025).

25. Ketata, M. A. et al. DiffDock-PP: Rigid Protein-Protein Docking with Diffusion Models. in International Conference on Learning Representations (ICLR) 1–10 (2023).

26. Proctor, E. A., Ding, F. & Dokholyan, N. V. Discrete molecular dynamics. Wiley Interdiscip. Rev. Comput. Mol. Sci. 1, 80–92 (2011).

27. Ho, J., Jain, A. & Abbeel, P. Denoising Diffusion Probabilistic Models. Adv. Neural Inf. Process. Syst. 33, 6840–6851 (2020).

28. Satorras, V. G., Hoogeboom, E. & Welling, M. E(n) Equivariant Graph Neural Networks. Proc. Mach. Learn. Res. 139, 9323–9332 (2021).

29. Gendron, P., Lemieux, S. & Major, F. Quantitative analysis of nucleic acid three-dimensional structures. J. Mol. Biol. 308, 919–936 (2001).

30. Battaglia, P. W. et al. Relational inductive biases, deep learning, and graph networks. arXiv preprint 1806.01261 1–40 (2018).

31. Wang, J., Zhang, D. Y., Budakoti, S. & Dokholyan, N. V. A Diffusion-Based Framework for Designing Molecules in Flexible Protein Pockets. Preprint at 10.1101/2025.05.27.656443 (2025).

32. Wang, J. & Dokholyan, N. V. Unified Protein-Small Molecule Graph Neural Networks for Binding Site Prediction. Preprint at 10.1101/2025.09.03.674017 (2025).

33. Joshi, C. K. et al. Grnade: Geometric Deep Learning for 3D Rna Inverse Design. 13th International Conference on Learning Representations, ICLR 2025 56390–56415 (2025).

34. Ding, F., Lavender, C. A., Weeks, K. M. & Dokholyan, N. V. Three-dimensional RNA structure refinement by hydroxyl radical probing. Nat. Methods 9, 603–608 (2012).

